# HIF1α is an essential regulator of steroidogenesis in the adrenal gland

**DOI:** 10.1101/2020.07.08.191783

**Authors:** Deepika Watts, Johanna Stein, Ana Meneses, Nicole Bechmann, Ales Neuwirth, Denise Kaden, Anja Krüger, Anupam Sinha, Vasileia Ismini Alexaki, Luis Gustavo Perez-Rivas, Stefan Kircher, Antoine Martinez, Marily Theodoropoulou, Graeme Eisenhofer, Mirko Peitzsch, Ali El-Armouche, Triantafyllos Chavakis, Ben Wielockx

## Abstract

Endogenous steroid hormones, especially glucocorticoids and mineralocorticoids, are essential for life regulating numerous physiological and pathological processes. These hormones derive from the adrenal cortex, and drastic or sustained changes in their circulatory levels affect multiple organ systems. Although a role for hypoxia pathway proteins (HPP) in steroidogenesis has been suggested, knowledge on the true impact of the HIFs (Hypoxia Inducible Factors) and oxygen sensors (HIF-prolyl hydroxylase domain-containing enzymes; PHDs) in the adrenocortical cells of vertebrates is scant. By creating a unique set of transgenic mouse lines, we reveal a prominent role for HIF1α in the synthesis of virtually all steroids under steady state conditions. Specifically, mice deficient in HIF1α in a part of the adrenocortical cells displayed enhanced levels of enzymes responsible for steroidogenesis and a cognate increase in circulatory steroid levels. These changes resulted in cytokine alterations and changes in the profile of circulatory mature hematopoietic cells. Conversely, HIF1α overexpression due to combined PHD2 and PHD3 deficiency in the adrenal cortex resulted in the opposite phenotype of insufficient steroid production due to impaired transcription of necessary enzymes. Based on these results, we propose HIF1α to be a central and vital regulator of steroidogenesis as its modulation in adrenocortical cells dramatically impacts hormone synthesis with systemic consequences. Additionally, these mice can have potential clinical significances as they may serve as essential tools to understand the pathophysiology of hormone modulations in a number of diseases associated with metabolic syndrome, auto-immunity or even cancer.

## Introduction

Steroidogenesis in the adrenal gland is a complex process of sequential enzymatic reactions that convert cholesterol into steroids, including mineralocorticoids and glucocorticoids (1). While glucocorticoids are regulated by the hypothalamic-pituitary-adrenal axis (HPA axis) and are essential for stress management and immune regulation (2, 3), aldosterone, the primary mineralocorticoid, regulates the balance of water and electrolytes in the body (4). As steroidogenesis is a tightly regulated process, proper control of adrenal cortex function relies on appropriate endocrine signaling, tissue integrity, and homeostasis (5). Accordingly, it has been suggested that inappropriately low pO_2_, or hypoxia, can lead to both structural changes in the adrenal cortex and interfere with hormone production (6-10).

Hypoxia inducible factors (HIFs) are the main transcription factors that are central to cellular adaptation to hypoxia in virtually all cells of our body. The machinery that directly controls HIF activity consists of the HIF-prolyl hydroxylase domain-containing enzymes (PHDs 1-3), which are oxygen sensors that hydroxylate two prolyl residues in the HIFα subunit under normoxic conditions, thereby marking the HIFs for proteasomal degradation. Conversely, oxygen insufficiency renders these PHDs inactive, leading to the binding of the HIF-complex to hypoxia responsive elements (HRE) in the promotor of multiple genes that ensure oxygen delivery and promote adaptive responses to hypoxia such as hematopoiesis, blood pressure regulation, and energy metabolism (reviewed in (11, 12)). Apart from directly activating hypoxia-responsive genes (13, 14), HIFs also indirectly influence gene expression by interfering with the activity of other transcription factors or systems. Of the most intensively studied HIFα genes, HIF1α has a ubiquitous pattern of expression in all tissues, whereas expression of the paralogue HIF2α is restricted to a selection of cell types (15, 16).

Recent *in vitro* and zebrafish studies have revealed a continuous cross talk between HIF and steroidogenesis pathways, along with potential interference in the production of aldosterone and glucocorticoids (17-20). There is also evidence suggesting a role for the hypoxia pathway in modulating glucocorticoid/glucocorticoid receptor (GR) signaling (21, 22). Importantly, these observations indicate a possible interplay of HIFs and PHDs in modulating the immune-regulatory actions of the HPA axis. Currently, there is huge interest in the development of HIF inhibitors and HIF stabilizers, and their influence on medicine is expected to become significant in the near future (23). However, as the role of HIFs/PHDs is both central and manifold with respect to maintaining oxygen homeostasis, a better understanding of the true impact of Hypoxia Pathway Proteins (HPPs) in the complex interplay of different essential physiological and pathological conditions, including in the adrenal cortex, assumes great importance.

We describe the creation and use of a unique collection of transgenic mouse lines that enabled an investigation of the role of HIFα subunits and PHDs in adrenocortical cells. Our results point towards a central role for HIF1α in the direct regulation of steroidogenesis in the adrenal gland and consequent changes in circulatory hormone levels. Importantly, chronic exposure of mice to such altered hormone levels eventually led to a dramatic decrease in essential inflammatory cytokines and profound dysregulation of circulatory immune cell profiles.

## Materials and Methods

### Mice

All mouse strains were housed under specific pathogen-free conditions at the Experimental Centre of the Medical Theoretical Center (MTZ, Technical University of Dresden – University Hospital Carl-Gustav Carus, Dresden, Germany). Experiments were performed with male and female mice aged between 8-16 weeks. No significant differences between the genders were observed. Akr1b7:cre-PHD2/HIF1^ff/ff^ (P2H1) or Akr1b7:cre-PHD2/PHD3^ff/ff^ (P2P3) lines were generated by crossing Akr1b7:cre mice (24) to PHD2^f/f^, HIF1α^f/f^ or PHD2^f/f^; PHD3^f/f^ as previously reported by us (25), and/or the reporter strain mTmG (26). All mice described in this report were born in normal Mendelian ratios. Mice were genotyped using primers described in supplementary Table 1. Histological analysis of the adrenal gland of Akr1b7:cre-mTmG^f/f^ reporter mice revealed zonal variation in the penetrance of cre-recombinase activity in the adrenal cortex of all individual mice (GFP^+^ staining). Peripheral blood was drawn from mice by retro-orbital sinus puncture using heparinized micro hematocrit capillaries (VWR, Darmstadt, Germany) and plasma separated and stored at −80 °C until further analysis. Mice were sacrificed by cervical dislocation and adrenals were isolated, snap frozen in liquid nitrogen, and stored at −80°C for hormone analysis or gene expression analysis. All mice were bred and maintained in accordance with facility guidelines on animal welfare and with protocols approved by the Landesdirektion Sachsen, Germany.

### Blood analysis

White blood cell counts were measured using a Sysmex automated blood cell counter (Sysmex XE-5000) (27).

### ACTH measurements

Plasma ACTH was determined using a radioimmunoassay, as per manufacturer’s instructions (ImmuChem Double Antibody hACTH 125 I RIA kit; MP Biomedicals Germany GmbH, Eschwege, Germany) (28).

### Hormone detection

Adrenal glands were incubated in disruption buffer (component of Invitrogen™ Paris™ Kit, AM 1921, ThermoFisher Scientific, Dreieich, Germany) for 15min at 4°C, homogenized in a tissue grinder, followed by incubation for 15 min on ice and further preparation. *Adrenal steroid hormones* were determined by LC-MS/MS as described elsewhere (29). *Catecholamines*, norepinephrine, epinephrine, and dopamine were measured by high pressure liquid chromatography (HPLC) coupled with electrochemical detection, as previously described (30).

### RNA extraction and qPCRs

RNA from adrenal glands and sorted cells was isolated using the RNA Easy Plus micro kit (Qiagen) (Cat. # 74034Qiagen). cDNA synthesis was performed using the iScript cDNA Synthesis Kit (BIO-RAD, Feldkirchen, Germany). Gene expression levels were determined by performing quantitative real-time PCR using the ‘Ssofast Evagreen Supermix’ (BIO-RAD, Feldkirchen, Germany). Sequences of primers used are provided in supplemental Table 2. Expression levels of genes were determined using the Real-Time PCR Detection System-CFX384 (BIO-RAD, Feldkirchen, Germany). All mRNA expression levels were calculated relative to β2M or EF2 housekeeping genes and were normalized using the ddCt method. Relative gene expression was calculated using the 2(-ddCt) method, where ddCT was calculated by subtracting the average WT dCT from dCT of all samples individually.

### Immunohistochemistry and immunofluorescence

For preparation of paraffin sections, adrenal glands were isolated, incubated in 4% formaldehyde at 4°C overnight, dehydrated, embedded in paraffin and cut into 5µm sections using a microtome. Sections were rehydrated and subjected to hematoxylin and eosin staining (H&E). For frozen sections, adrenal glands were embedded in O.C.T Tissue-Tek (A. Hartenstein GmbH, Würzburg, Germany) and stored at −20°C. For H&E staining of frozen sections (7µm), samples were first fixed in cold acetone before staining. For immunofluorescence, sections were fixed in cold acetone, air-dried, washed with phosphate-buffered saline containing 0.1% Tween-20, blocked with 5% normal goat serum followed by primary antibody staining (CD31/PECAM – 1:500 (31)) or GFP Polyclonal (Antibody ThermoFischer Scientific – 1:200) overnight at 4°C and subsequent secondary antibody staining. After counterstaining with DAPI, slides were mounted in fluorescent mounting medium and stored at 4 °C until analysis.

### Microscopy

Both brightfield and fluorescent images were acquired on an ApoTome II Colibri (Carl Zeiss, Jena, Germany). Images were analyzed using either Zen software (Carl Zeiss, Jena, Germany) or Fiji (ImageJ distribution 1.52K). Fiji was used to quantify lipid droplet sizes and CD31 staining.

### Meso Scale Discovery

Meso Scale Discovery (MSD, Rockville, Maryland) was used to measure cytokines in plasma samples using the MSD plate reader (QuickPlex SQ 120). Cytokine concentrations were calculated by converting the measured MSD signal to pg/ml using a standard curve. All values below that of blank (control) were considered as zero. Finally, all cytokine concentrations in individual P2H1 mice were normalized to the average value of WTs for every independent experiment; and the average WT value was set as 1.

### Next generation sequencing

For RNAseq analysis, adrenal glands from Akr1b7:cre-PHD2/HIF1/mTmG^fff/fff^ and Akr1b7:cre-mTmG^f/f^ (control) mice were isolated directly into the lysis buffer of the RNeasy Plus Micro Kit, RNA was isolated according to manufacturer’s instructions, and SmartSeq2 sequencing was performed (SmartSeq2 and data analysis in Supplemental Data). Flow cytometry and cell sorting were performed as described previously (32).

### Read Quantification

Kallisto v0.43 was first used to generate an index file from the transcript file, which can be downloaded from :ftp://ftp.ebi.ac.uk/pub/databases/gencode/Gencode_mouse/release_M12/gencode.vM12.transcripts.fa.gz. Kallisto v0.43 was then run on all the fastq files using parameters “quant --single -l 75 – s 5 -b 100” to quantify reads for the genes.

### Differential Gene Expression Quantification

Complete cDNA sleuth v0.30.0 (an R package) was used to evaluate differential expression. The command “sleuth_prep” was run with parameter “gene_mode=TRUE”. Two separate error models were fit using “sleuth_fit” wherein the first was a “full” model with gender and experimental condition as covariates, while the second was a “reduced” model with only gender as the covariate. “sleuth_lrt” (Likelihood Ratio Test) was used to evaluate differential gene expression by comparing the full model and the reduced model.

### Statistical analyses

All data are presented as mean ± SEM. Data (WT control versus transgenic line) were analyzed using the Mann–Whitney U-test, unpaired t-test with Welch’s correction as appropriate (after testing for normality with the F test) or as indicated in the text. All statistical analyses were performed using GraphPad Prism v7.02 for Windows (GraphPad Software, La Jolla California USA, www.graphpad.com). Significance was set at p<0.05; “n” in the figure legends denotes individual samples.

## Results

### A new mouse model to study the effects of alterations in hypoxia pathway proteins (HPPs) in the adrenal cortex

We took advantage of the adrenal cortex-specific Akr1b7:cre recombinase mouse line (25) to investigate the effects of adrenocortical HPPs on the structure and functions of the adrenal gland. When combined with the mTmG reporter strain (26), we show up to 40% targeting among all cortical cells (Figure 1A). Next, we generated the Akr1b7:cre-PHD2/HIF1^ff/ff^ mouse line (henceforth designated P2H1) by combining Akr1b7:cre mice with PHD2 and HIF1α floxed mice (24). Genomic PCRs on DNA and qPCR analysis using mRNA from whole adrenal glands revealed targeting of *PHD2* and *HIF1α*, when compared to WT littermates (Figure 1B-C). Importantly, in P2H1 mice, we even detected a significant increase in *HIF2α* mRNA but not of *PHD3*, which is in line with our earlier report of enhanced HIF2α-activity in PHD2/HIF1α-deficient cells (24). Therefore, we explored the expression profile of a number of downstream genes known to be transactivated by HIF2α (33-35) and found a significant increase in *Vegfa, Hmox1*, and to a lesser extent *Bnip3* levels, underscoring the functionality of the P2H1 mouse line (Figure 1E).

**Figure 1.**
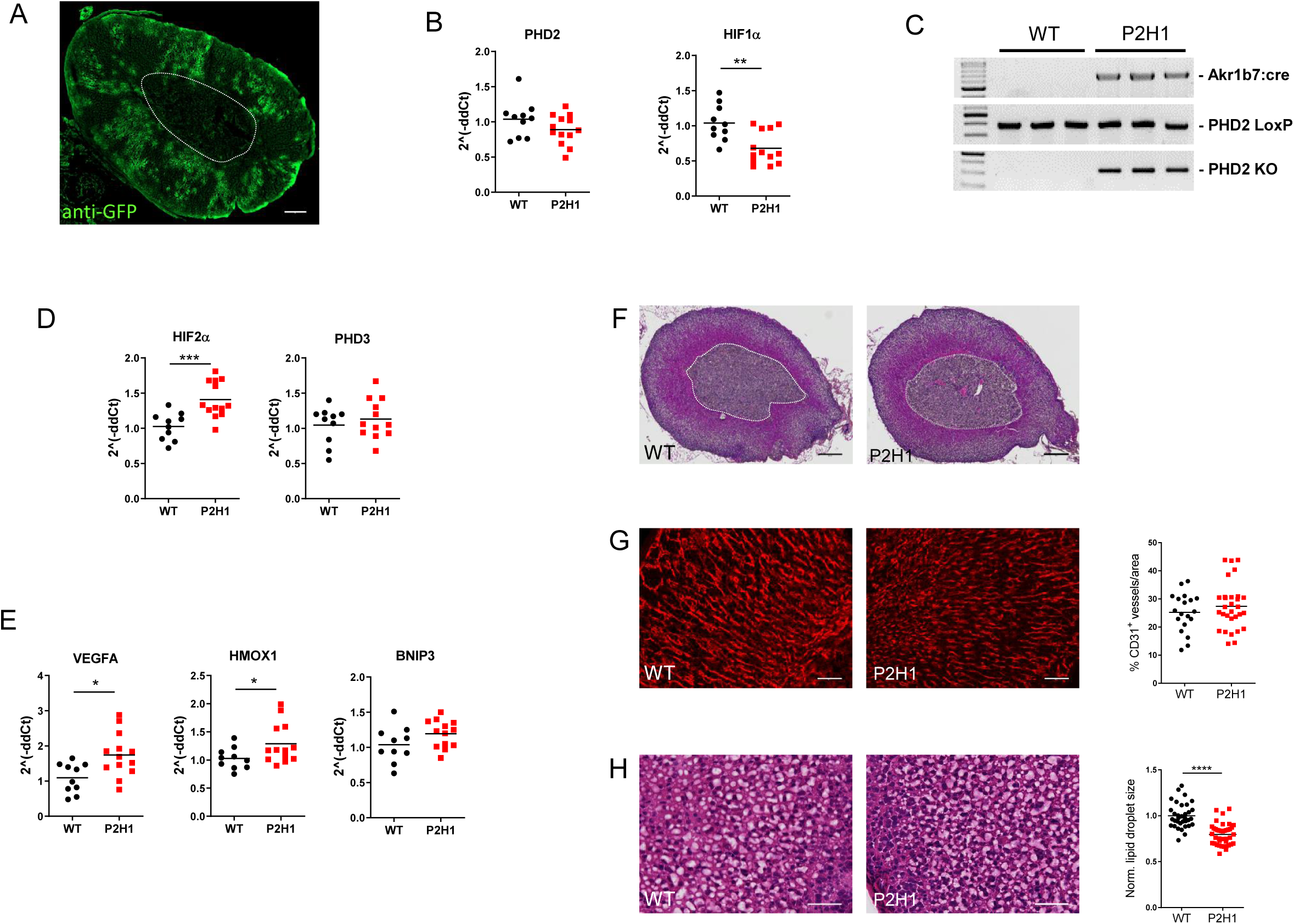
Characterization of the Akr1b7:cre-P2H1^ff/ff^ mouse line with cortex-specific targeting of hypoxia pathway proteins. A: Representative immunofluorescent image of anti-GFP stained (GFP+) area in the adrenal cortex of the Akr1b7:cre-mTmG mouse line. Region enclosed within the white dotted line represents the medulla and it demarcates the medulla from the cortex (scale bar, 100 μm). B: qPCR-based mRNA expression analysis of PHD2 and HIF1α in entire adrenal tissue from P2H1 mice and WT littermates (n=10-13). Relative gene expression was calculated using the 2^(^−^ddCt) method. The graphs represent data from 2 independent experiments. C: Genomic PCRs for Akr1b7:cre (650bp), PHD2 LoxP (400bp), and PHD2 KO (350bp) in DNA derived from whole adrenal glands of WT and P2H1 mice. D-E: Relative gene expression analysis using mRNA from the entire adrenal tissue in P2H1 mice and their WT counterparts (n=10-13). All graphs represent data from 2 independent experiments. F: Representative images (magnification 20x) of paraffin sections of adrenal glands (H&E) from 8-week old WT and P2H1 mice (scale bars represent 100μm). G: Representative immunofluorescent images of CD31^+^ endothelial cell staining in adrenal gland sections from WT and P2H1 mice (scale bars represent 50μm). Graph in the right-side panel represents quantification of CD31^+^ area as a fraction of total tissue area. Each data point represents a single measurement of the cortical area in the adrenal gland (collection of n=6 vs 11 individual mice). H: Representative images of cryo-sections of WT and P2H1 adrenal glands (H&E) (scale bars represent 50μm). Graph in the right-side panel represents the normalized average size of an individual lipid droplet per section of adrenal gland tissue in WT versus P2H1 mice. Measurements were made from 6 sections per mouse. (n=8 individual adrenals per genotype). The graphs in panels G and H are representative of 2 independent experiments. Statistical significance was defined using the Mann-Whitney U test (*p<0.05; **p<0.005; ***p<0.001; ****p<0.0001).

### Morphological changes in the adrenal cortex of P2H1 mice

To evaluate the impact of changes in HIF1α and/or HIF2α activity in adrenocortical cells, we analyzed adrenal gland morphology using H&E staining on paraffin sections but found no differences between P2H1 mice and WT littermates in the structure of the adrenal gland, especially, at the side of the cortex of P2H1 mice in comparison to WT littermates (Figure 1F). As we detected a significant increase in *Vegfa* in the adrenal glands of P2H1 mice, we used CD31 staining to quantify endothelial cells but detected no significant differences between P2H1 and WT mice (Figure 1G). Remarkably, H&E staining on cryosections of P2H1 adrenal glands revealed significantly smaller lipid droplets in the adrenocortical cells (Figure 1H), an effect that is reported to be correlated with greater conversion of cholesterol into pregnenolone (10).

### Modulation of HPPs in the adrenal cortex enhances synthesis and circulatory levels of steroid hormones

Next, to verify if the observed changes in lipid droplets indeed led to changes in steroidogenesis, we quantified steroid hormones and their precursor levels by LC-MS/MS in the adrenal gland and in plasma. Quantification revealed a significant increase in virtually all of the hormones tested in P2H1 adrenal glands compared to WT littermates (Figure 2A), and importantly, a corresponding increase of progesterone, corticosterone, and aldosterone was found in the plasma (Figure 2B). These observations clearly indicate that central HPPs have an impact on steroidogenesis in the murine adrenal gland and on circulatory levels of steroid hormones.

**Figure 2:**
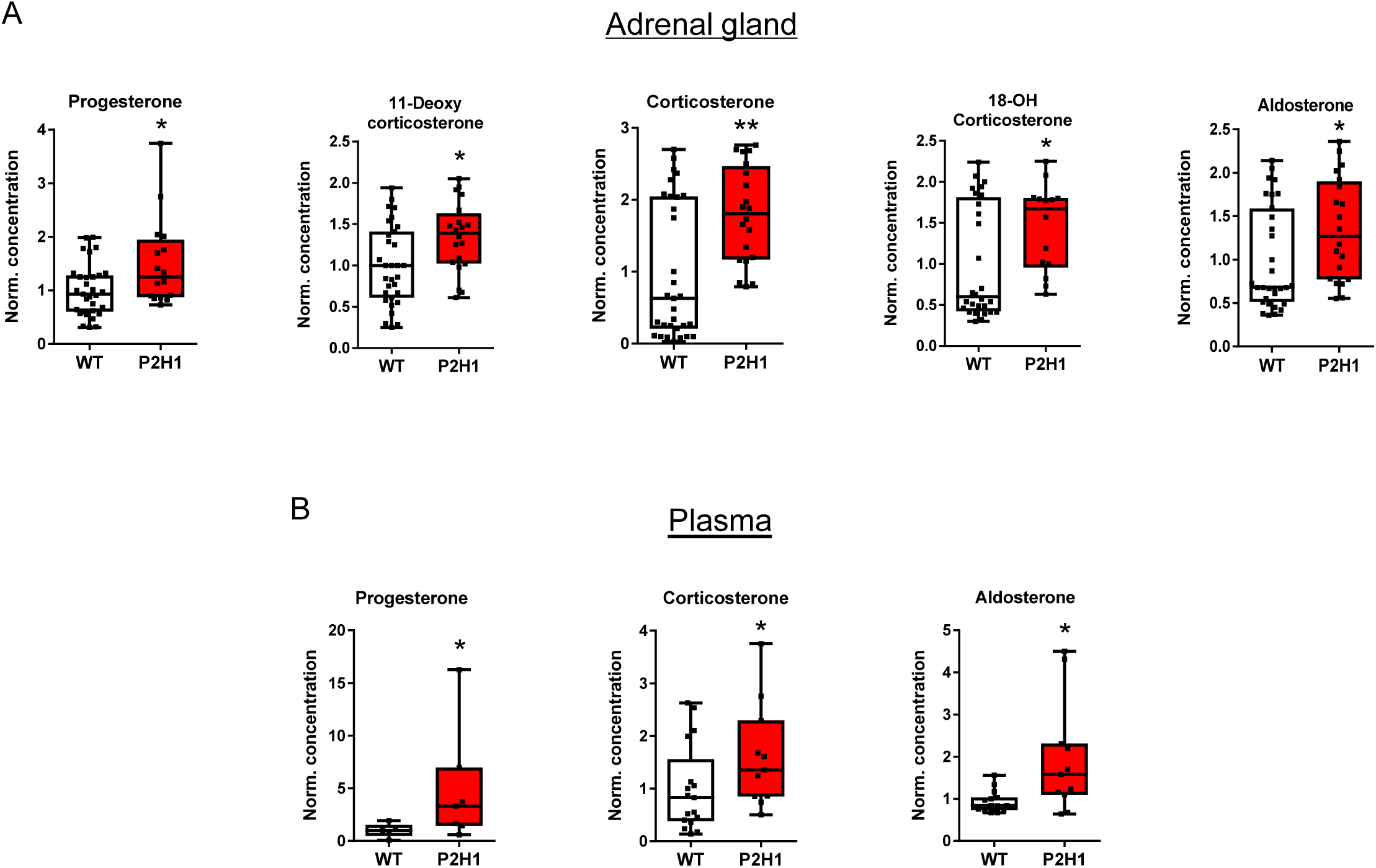
Adrenal cortex-specific loss of PHD2 and HIF1 leads to enhanced steroidogenesis in P2H1 mice. A: Box and whisker plots showing steroid hormone measurements in adrenal glands from WT mice and compared to littermate P2H1 mice (n=20-31 individual adrenal glands). B: Box and whisker plots showing steroid hormone measurements in the plasma of individual mice (n=5-17). All data were normalized to average measurements in WT mice. The graphs are a representative result of at least 3 independent experiments. Statistical significance was defined using the Mann-Whitney U test (*p<0.05; **p<0.005).

### Downstream effects of the chronic increase in the steroidogenesis

Previous reports have stated that glucocorticoids can regulate catecholamine production in the adrenal medulla (36, 37); therefore, we also measured dopamine, norepinephrine, and epinephrine levels in the samples used to quantify steroid levels (as above). However, we found no difference between P2H1 and WT littermates in any of the catecholamines quantified (Supplementary Figure 1A). Further, although increased steroid levels often result in a negative feedback loop affecting ACTH secretion from the pituitary (38), P2H1 mice displayed no such differences compared to WT littermates (Supplementary Figure 1B), nor did they have any difference in serum potassium levels or blood glucose levels (Supplementary Figure 1C-D). Taken together, in contrast to the systemic effects induced by acute and high levels of circulatory cortical hormones (e.g. corticosterone, aldosterone) (3, 4), the P2H1 mice display moderate but chronically enhanced levels of cortical hormones at the described time points.

### Loss of PHD2/HIF1α in adrenocortical cells impacts gene expression related to steroidogenesis

Previous *in vitro* studies and reports on HIF1α alterations in zebrafish larvae have suggested negative regulation of StAR, the mitochondrial cholesterol transporter (7, 17, 20). However, data on the effects of HPP alterations in adrenal cortex of mice is scant at best. Therefore, to assess the impact of HIF1α-deletion and/or HIF2α-upregulation in adrenal cortical cells, we performed broad transcription analysis of proteins/enzymes involved in steroidogenesis using mRNA from whole adrenals. Our results reveal that almost all of the gene products tested showed either a significant increase or a tendency to do so, including key enzymes like *StAR, Cyp11a1, Cyp21a1* and *Cyp11b1* (Figure 3A).

**Figure 3:**
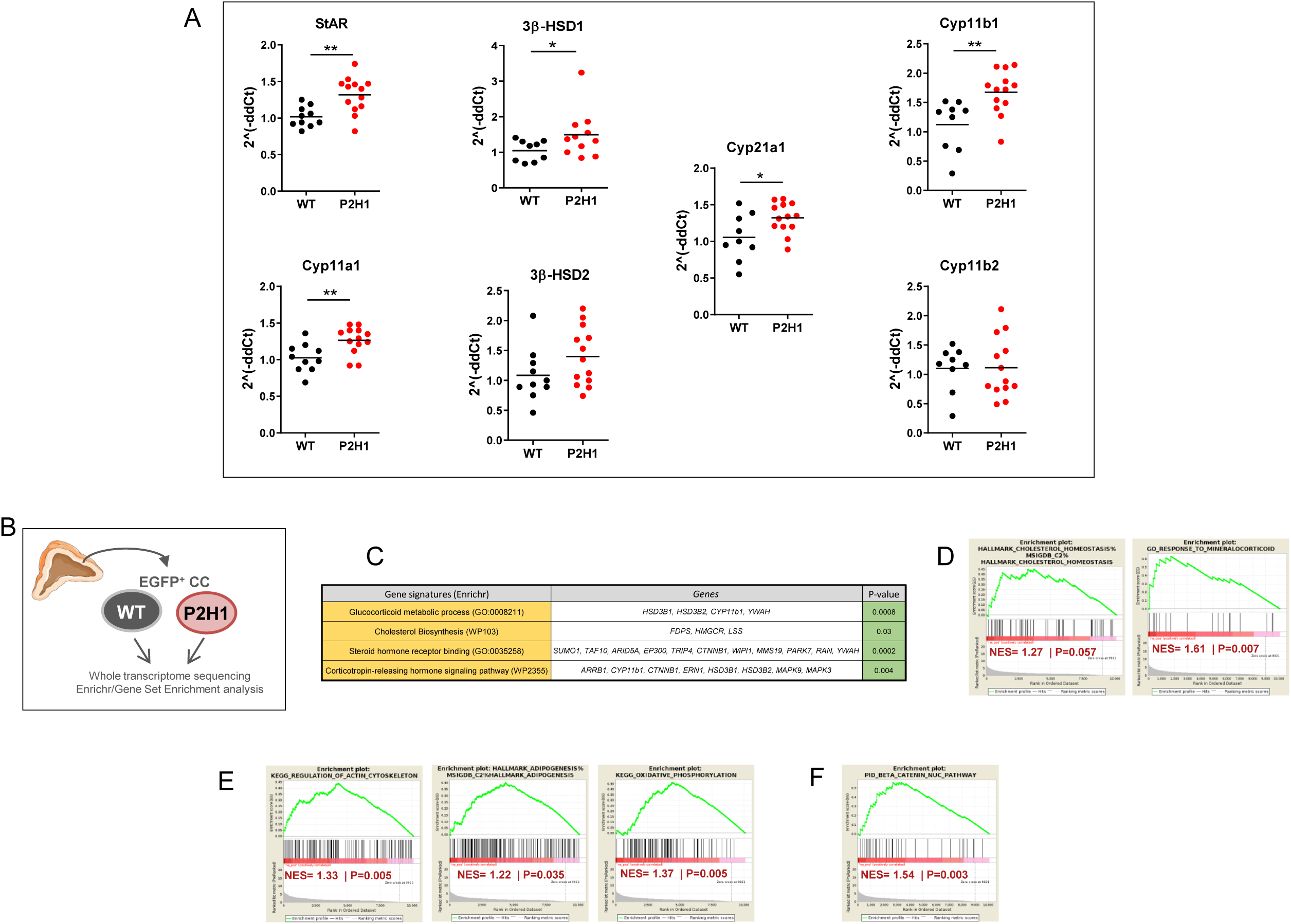
Gene expression analysis of P2H1 adrenocortical cells. A: Gene expression analysis of enzymes involved in the steroidogenesis pathway using mRNA from whole adrenals from P2H1 mice and WT counterparts (n=10-13). All graphs are the result of 2 independent experiments. Statistical significance was defined using the Mann-Whitney U test (*p<0.05; **p<0.005). B: Schematic overview of the RNAseq approach which compared sorted GFP^+^ cells from WT controls and P2H1 mice (n=3). C: Gene signature analysis using Enrichr. D. Gene set enrichment analyses (GSEA) showed positive signatures for steroidogenesis related pathways. E: prominent HIF-related pathways. F: the β-catenin nuclear pathway.

To further characterize this phenotype driven by the HPPs, we performed *next generation sequencing* (NGS) and compared the steady state transcriptomes of P2H1 and WT littermate mice (Figure 3B). For this, we specifically created the Akr1b7:cre-PHD2/HIF1/mTmG^fff/fff^ mouse line (P2H1 reporter mice) to study only targeted adrenal cortex cells, with Akr1b7:cre-mTmG^f/f^ animals used as controls. Bulk RNAseq was performed on GFP^+^-sorted adrenal gland cells as described previously (39) and gene signatures of the various lineages were evaluated using Enrichr or gene set enrichment analyses (GSEA). Concurring with the previous results, we found a number of significant signatures related to the process of steroid synthesis in adrenocortical cells or their response to it (Figure 3C-D). Notably, GSEA also revealed known HIF-dependent associations including, actin cytoskeleton (40, 41), adipogenesis (42) and oxidative phosphorylation (43) (Figure 3E). Furthermore, P2H1 cortical cells also displayed a positive signature related to the regulation of nuclear β-catenin signaling, which is known to be primarily activated in the zona glomerulosa with potential hyperplasic effects (44) (Figure 3F).

### Modulated adrenocortical HPPs skew cytokine production and leukocyte numbers

As several studies have reiterated a crucial role for glucocorticoids in immunomodulation (3, 45), and Cushing’s syndrome has been described to be accompanied by immune deficiency (3, 38, 46), we measured circulatory cytokine levels. We report a substantial overall decrease in the levels of both pro- and anti-inflammatory cytokines, with the exception of the chemokine and neutrophil attractant CXCL1, which increased almost 2-fold (Figure 4A). Glucocorticoids have been repeatedly shown to promote apoptosis-mediated reduction of lymphocytes (47) and eosinophil reduction (48), along with neutrophilia due to enhanced recruitment from the bone marrow (49). Therefore, we enumerated the various white blood cell (WBC) fractions in P2H1 mice and compared it with that of their WT littermates, which revealed a significant reduction in both lymphocyte and eosinophil fractions (Figure 4B) accompanied by marked elevation in neutrophils (>70% compared to WT) (Figure 4C). Taken together, our data reveal a critical role for HPPs in steady-state cytokine levels and leukocyte numbers, probably through alterations in steroidogenesis pathways.

**Figure 4:**
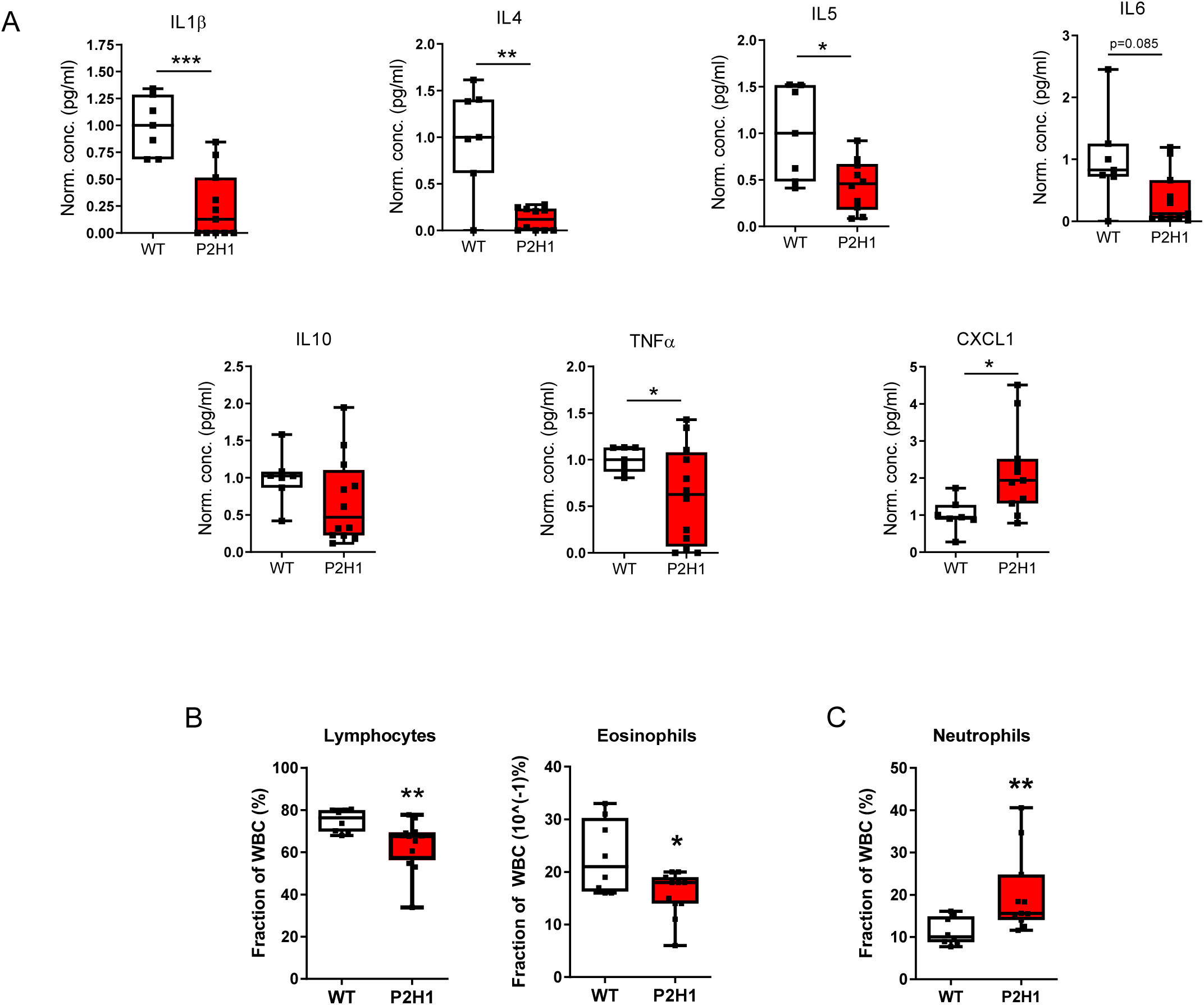
Immune system changes in P2H1 mice. A: Box and whisker plots representing levels of pro/anti-inflammatory cytokines measured in the plasma of P2H1 mice and WT littermate controls (n=7-12). All data were normalized to the average value seen in WT mice. Each dot represents data from one animal. B: Box and whisker plots showing percentage lymphocytes and eosinophils in circulation which revealed reduced fractions in P2H1 mice compared to WT controls. C: Greater numbers of circulating neutrophils in P2H1 mice compared to WT littermates. All graphs represent pooled results of 2 independent experiments. Statistical significance for cytokines in panels A and B was defined using the Mann-Whitney U test, except for TNFα, where the Unpaired t test with Welch’s correction was used after verifying data normality. (*p<0.05; **p<0.005; ***p<0.001).

### HIF1α inversely regulates steroidogenesis

To extend our understanding of the role of HIF1α and/or HIF2α in adrenocortical cells, we created the Akr1b7:cre-PHD2/PHD3^ff/ff^ mouse line (designated as P2P3), which showed adequate activation efficiency upon genomic PCRs of whole adrenal tissue (supplementary Figure 2). Intriguingly and in contrast to hormone levels in the adrenal glands of the P2H1 mice, P2P3 adrenal glands displayed a marked decrease in corticosterone and aldosterone levels, along with a cognate reduction in their precursors, both in the adrenal gland (Figure 5A) and in circulation (Figure 5B). These results clearly suggest that steroidogenesis is dependent on HIF1α but not HIF2α. To further confirm this observation, we performed mRNA expression analyses to identify the levels of central enzymes, similar to that performed in P2H1 mice, and demonstrate an overall decrease in these enzymes (Figure 6A). This observation is contrary to that seen in the P2H1 mice but fits neatly with the observed reduction in steroid levels in the P2P3 mice, thereby underscoring the central role of HIF1α (Figure 6B).

**Figure 5:**
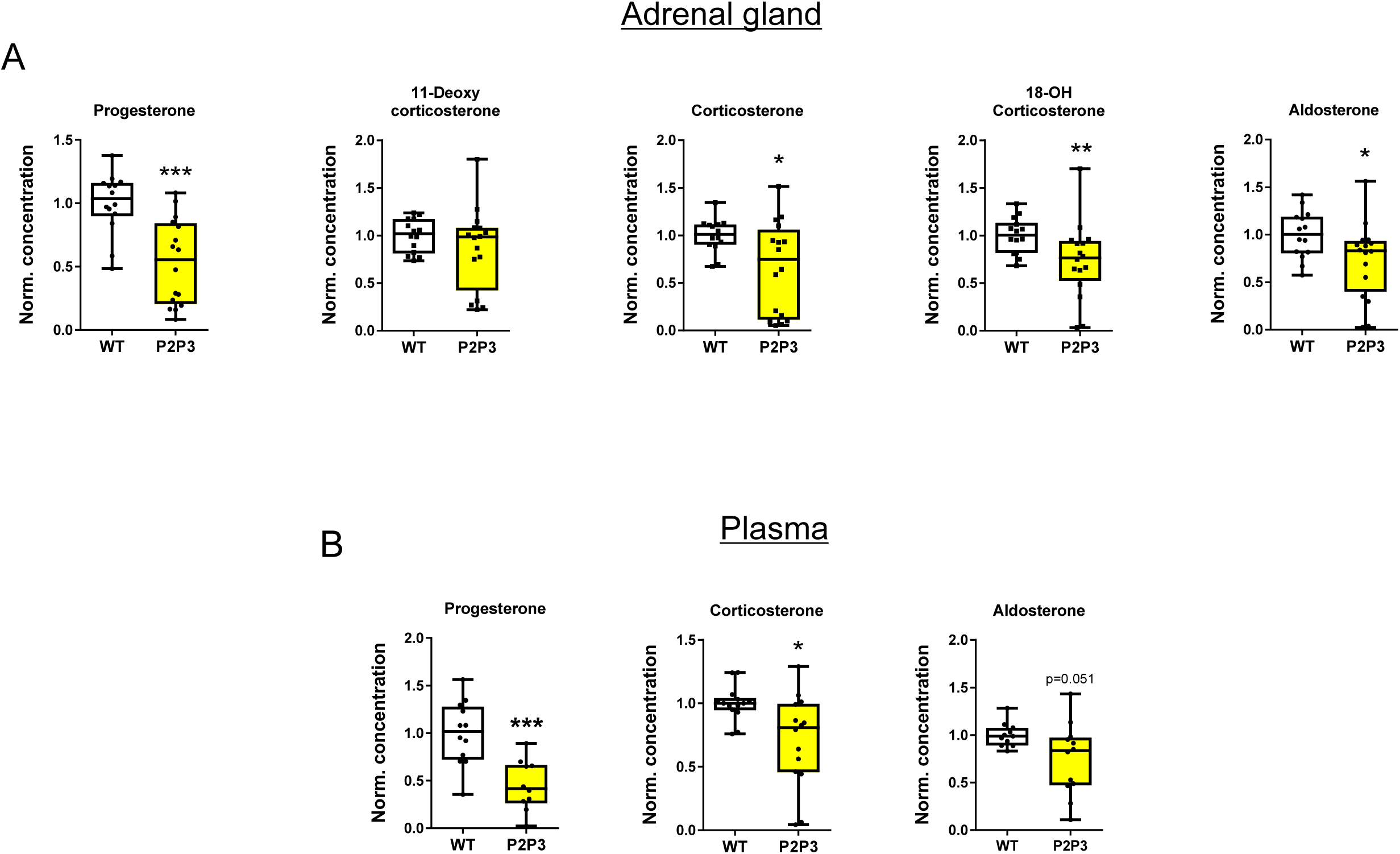
Adrenal cortex-specific loss of PHD2 and PHD3 leads to reduced steroidogenesis in mice. A: Box and whisker plots showing steroid hormone levels in the adrenal glands of WT mice and compared to that of littermate P2H1 mice (n=14-16 individual adrenal glands). B: Box and whisker plots showing steroid hormone measurements in the plasma of individual mice (n=10-12). All data were normalized to the average value of WT mice and graphs are representative of at least 3 independent experiments. Statistical significance was defined using the Mann-Whitney U test for progesterone, 11-deoxycorticosterone, and 18-OH corticosterone. Unpaired t test with Welch’s correction was used for corticosterone and aldosterone after verification of data normality (*p<0.05; **p<0.005; ***p<0.001).

**Figure 6:**
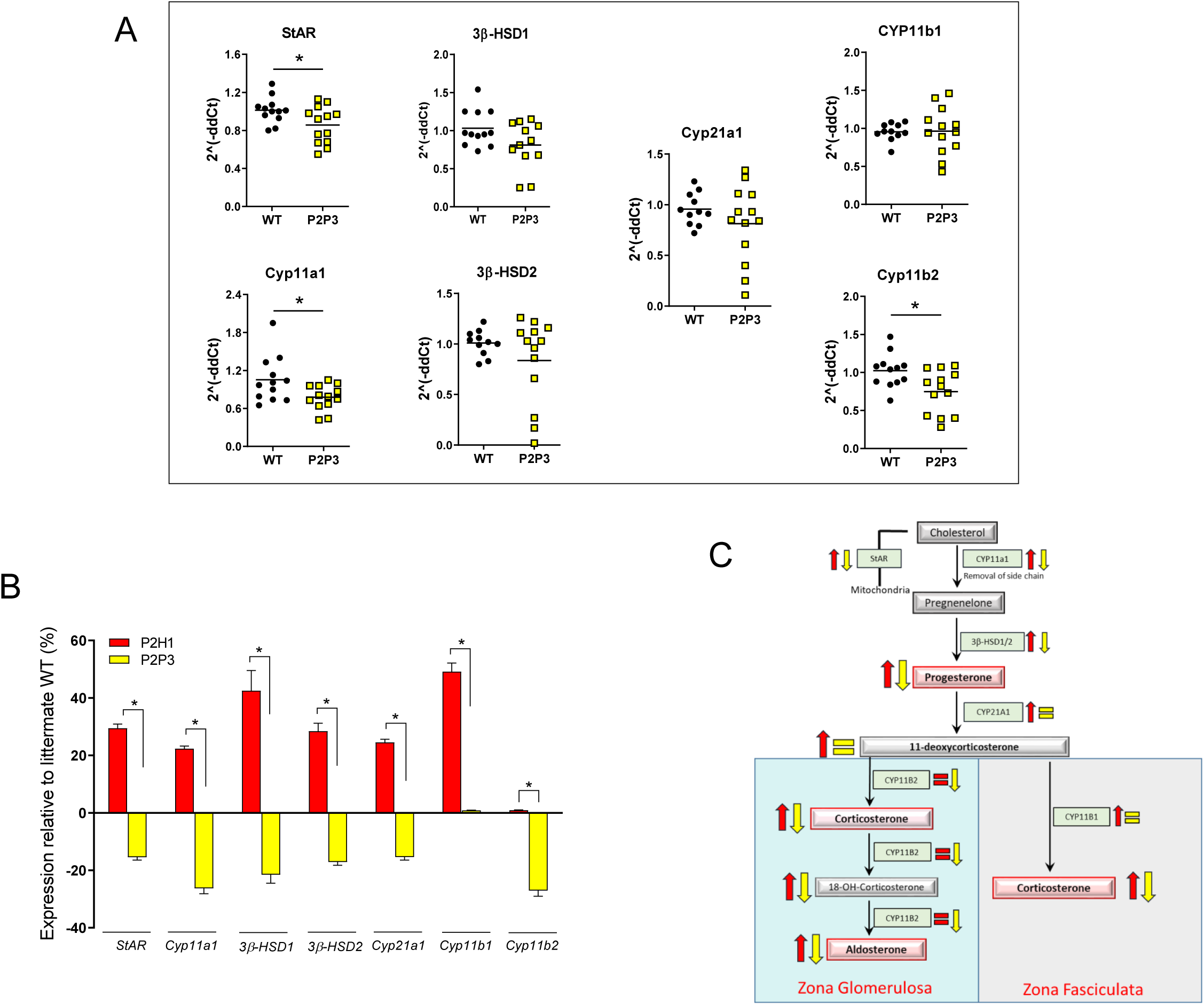
Inverse regulation of steroidogenesis in P2P3 mice compared to P2H1 mice. A: Gene expression analysis of enzymes involved in the steroidogenesis pathway in P2P3 mice and their WT counterparts (n=12-13) was performed in mRNA from entire adrenal glands. All graphs represent pooled data from at least 3 independent experiments. Statistical significance was defined using the Mann-Whitney U test (*p<0.05). B: Relative expression profile of all genes analyzed from the adrenal glands of P2H1 and P2P3 mice and compared to their respective WT littermates. Statistical significance was defined using an unpaired multiple t-test (n=13; Benjamini, Krieger and Yekutieli method; *p<0.0001 for all individual genes). C: schematic overview of all changes in adrenocortical enzymes and their corresponding hormones and intermediates reported here in P2H1 (red) and P2P3 (yellow) mice.

## Discussion

Here, by using a unique collection of adrenocortical-specific transgenic mouse lines, we identify HIF1α as a central transcription factor that regulates the steroidogenesis pathway by regulating key enzymes. Notably, this directly modifies the entire spectrum of steroid hormones, both in the adrenal gland and in circulation, which eventually impacts the availability of a variety of cytokines.

Studies on the role of HIFs in the regulation of steroidogenesis *in vitro* are few, apart from those in zebra fish larvae that describe differential regulation of the enzymes involved in the steroid pathway (7, 18, 20). However, to the best of our knowledge, there are no mouse models to study the role of HPPs in adrenal cortical cells. Undoubtedly, such models would help us to better understand the crosstalk between HPPs and adrenal steroid metabolism, while simultaneously serving as an essential tool to study the pathophysiology of multiple conditions associated with dramatically altered steroid hormone levels (2). Ablation of HIF1α revealed an important role for this transcription factor in steroidogenesis, which concurs with results from previous studies (20, 50). However, our findings that HIF1α deletion results in the upregulation of mRNA of a vast majority of steroid-related enzymes is counterintuitive to the nature of this transcription factor (12, 51), and therefore we believe this effect is most likely indirect with potential involvement of one or more transcriptional repressors (13, 52, 53). This type of transcriptional regulation of adrenal steroidogenesis has already been suggested with miRNAs, which are endogenous noncoding single-stranded small RNAs that suppress the expression of various target genes (54). Hu and colleagues have demonstrated that a HIF1α-dependent miRNA, miRNA-132, attenuates steroidogenesis by reducing StAR protein levels (55), and similar mechanisms have reported for *Cyp11B2* via miR-193a-3p (56, 57), and *Cyp11B1* and *Cyp11B2* via miR-10b (8). Thus, these new mouse lines will be of great value for in-depth studies on the complex background of HIF1α involvement in the expression patterns of steroidogenesis-related miRs.

Our RNAseq analysis of Akr1b7^+^ P2H1 adrenocortical cells not only unearthed several genetic signatures directly associated with steroidogenesis, but a number of GSEAs revealed prominent HIF-dependent phenotypes previously identified in a variety of other cell types. Recently, we have described a significant role for HIF2α in the regulation of the actin cytoskeleton, especially in facilitating enhanced neutrophil migration through very confined environments (41), HIF1α has also been associated with cytoskeleton structure and functionality in a number of cell lineages (reviewed in (40)); this is apart from its role in energy metabolism wherein enhanced oxidative phosphorylation has been demonstrated in various HIF1α-deficient cell lineages (43). Therefore, it will be of interest to further explore changes in multiple metabolites that are directly or indirectly-associated with the TCA cycle to find a potential link with the overall changes described here.

Glucocorticoids and aldosterone are both essential for homeostasis and their substantial increase in P2H1 mice was intriguing, given their pivotal role in immune suppression (3, 58) and blood pressure regulation, respectively. Previous studies have shown that aldosterone not only increases the expression of the potassium channels that secrete potassium but also stimulates K-absorptive pumps in the renal cortex and medulla, thereby stabilizing and maintaining renal potassium excretion (59), a situation we also observed in the P2H1 mice. The significant increase in glucocorticoids upon HIF1α deletion was clearly associated with immunosuppression, as demonstrated by an overall decrease in both pro- and anti-inflammatory cytokines in circulation, and these observations mirror other reports of immune modulation due to enhanced glucocorticoid levels. Such glucocorticoid elevation can eventually even result in dramatic immune deficiency, for example, as seen in Cushing’s disease (3, 38, 45, 58).

Intriguingly, we found serum CXCL1 to be significantly enhanced in P2H1 mice, probably because as a central neutrophil attractant it was associated with the massive increase in circulatory neutrophils seen in these mice. It is known that enhanced neutrophil recruitment from the bone marrow is directly associated with glucocorticoids (49), as is their overall survival (60, 61).

An essential role of HIF1α, but not HIF2α, in the modulation of enzymes and adrenocortical hormones could be further corroborated by the contrasting results seen in the P2P3 mice. Specifically, compared to P2H1 mice, the expression profile of virtually all steroidogenesis regulating enzymes was dramatically inverted in the P2P3 mice, which resulted in an overall impairment of the steroidogenesis pathway. Therefore, these mouse lines will also be helpful to study the potential impact of dramatically modulated steroid levels in a variety of clinically relevant diseases including metabolic and auto-immune disorders.

In summary, we reveal a prominent role for HIF1α as a central regulator of steroidogenesis in mice as two distinct transgenic mouse lines showed persistent but contrasting changes in corticosterone and aldosterone concentrations at levels sufficient to modulate systemic cytokine levels and leukocyte numbers. These P2H1 and P2P3 mouse strains are of significant importance in further exploring the impact of HIF1α in adrenocortical cells and as an essential component in regulation of steroidogenesis-mediated systemic effects.

## Acknowledgments

This work was supported by grants from the DFG (German Research Foundation) within the CRC/Transregio 205/1, Project No. 314061271 – TRR205, “The Adrenal: Central Relay in Health and Disease” (A02) to B.W., T.C, A-E-A.; B.W. was supported by the Heisenberg program, DFG, Germany; WI3291/5-1 and 12-1). We would like to thank Dr. Vasuprada Iyengar for English Language and content editing.

## Conflict-of-interest

The authors have declared that no conflict of interest exists.

## Author contributions

D.W. designed and performed the majority of experiments, analysed data, and contributed in writing the manuscript. J.S., D.K., A.K., performed experiments and analysed data. A.Me. designed several mouse lines and contributed to the discussion. N.B., A.N., A.E.A. and T.C. provided tools and contributed to the discussion. G.E. and M.P. provided tools, analyzed data and contributed to the discussions. V.I.A. contributed to the discussions. A.Ma. provided essential tools. A.S. performed deep sequencing analysis. L.G.P-R. and M.T. performed ACTH measurements and contributed to the discussion. B.W. designed and supervised the overall study, analysed data, and wrote the manuscript.

---

**Supplementary figure 1:**
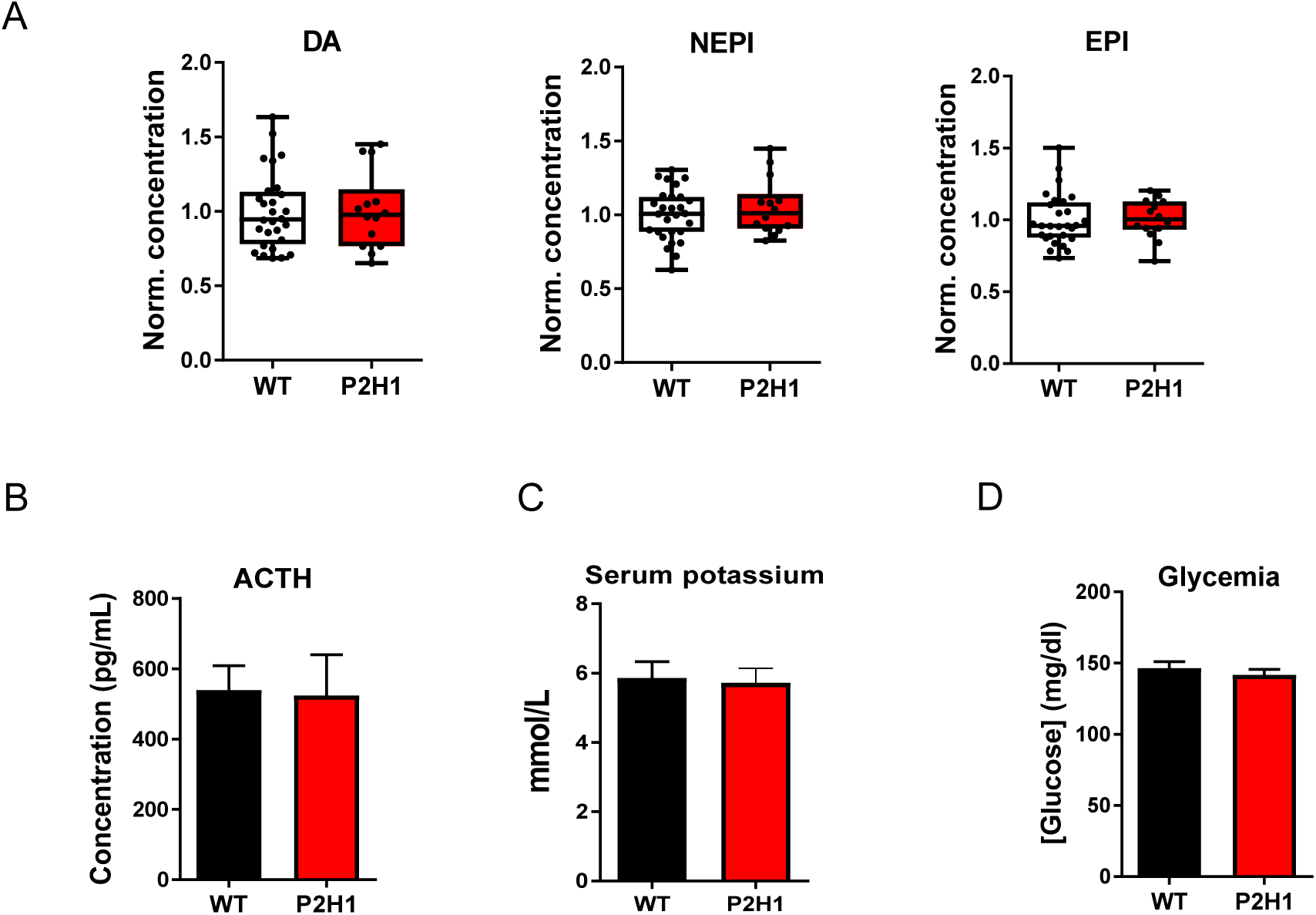
Downstream effects of increased steroidogenesis. A. Box and whisker plots showing normalized concentrations of all catecholamines (dopamine, norepinephrine (NEPI), and epinephrine (EPI) measured in entire adrenal glands of P2H1 mice and their WT counterparts (n=14-28). Bar graphs represent, respectively, B. Plasma ACTH concentration (n=7-14), C. potassium levels in the serum of WT vs P2H1 mice (n=9-11), D. Blood glucose levels in P2H1 mice vs WT littermate controls (n=5-8). Statistical significance was defined using the Mann-Whitney U test.

**Supplementary figure 2:**
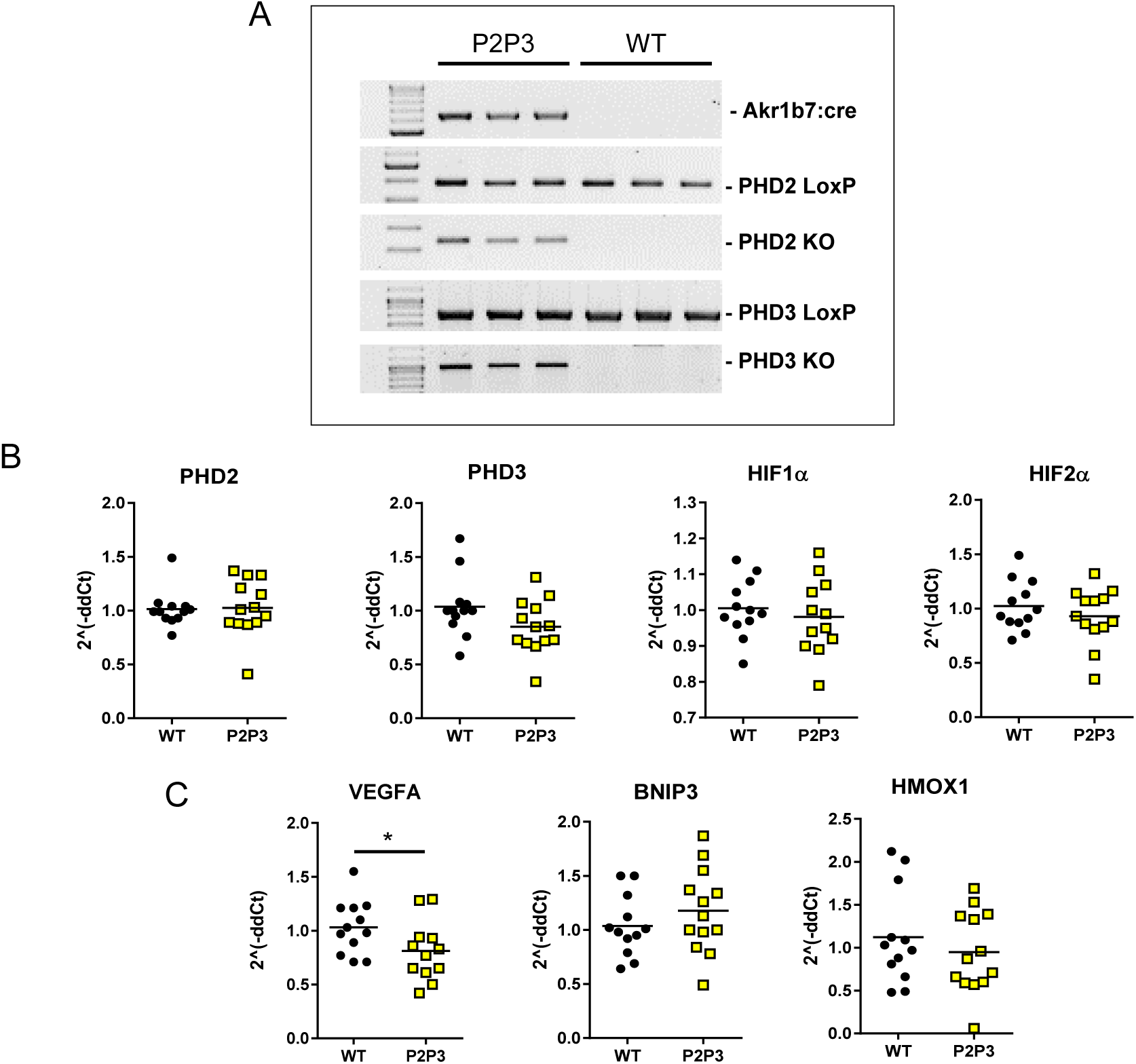
Genetic identification of the Akr1b7:cre-P2P3 strain. A. Genomic PCRs for Akr1b7:cre, PHD2 LoxP (400bp), PHD2 KO (350 bp), PHD3 LoxP (840bp) and PHD2 KO (1000bp) in entire adrenal gland tissue from P2P3 mice and their WT counterparts. B. Relative gene expression analysis by qPCR for *PHD2, PHD3, HIF1α* and *HIF2 α* in mRNA from entire adrenal glands of P2P3 and WT counterparts (n=12-13) C. qPCR as in panel B, but for *VEGFA, HMOX1, BNIP3* **(E)**. Relative gene expression was calculated using the 2^(^−^ddCt) method. The graphs are a representative result of 3 independent experiments. Statistical significance was defined using the Mann-Whitney U test – one tailed (*p<0.05).

**Table I:**
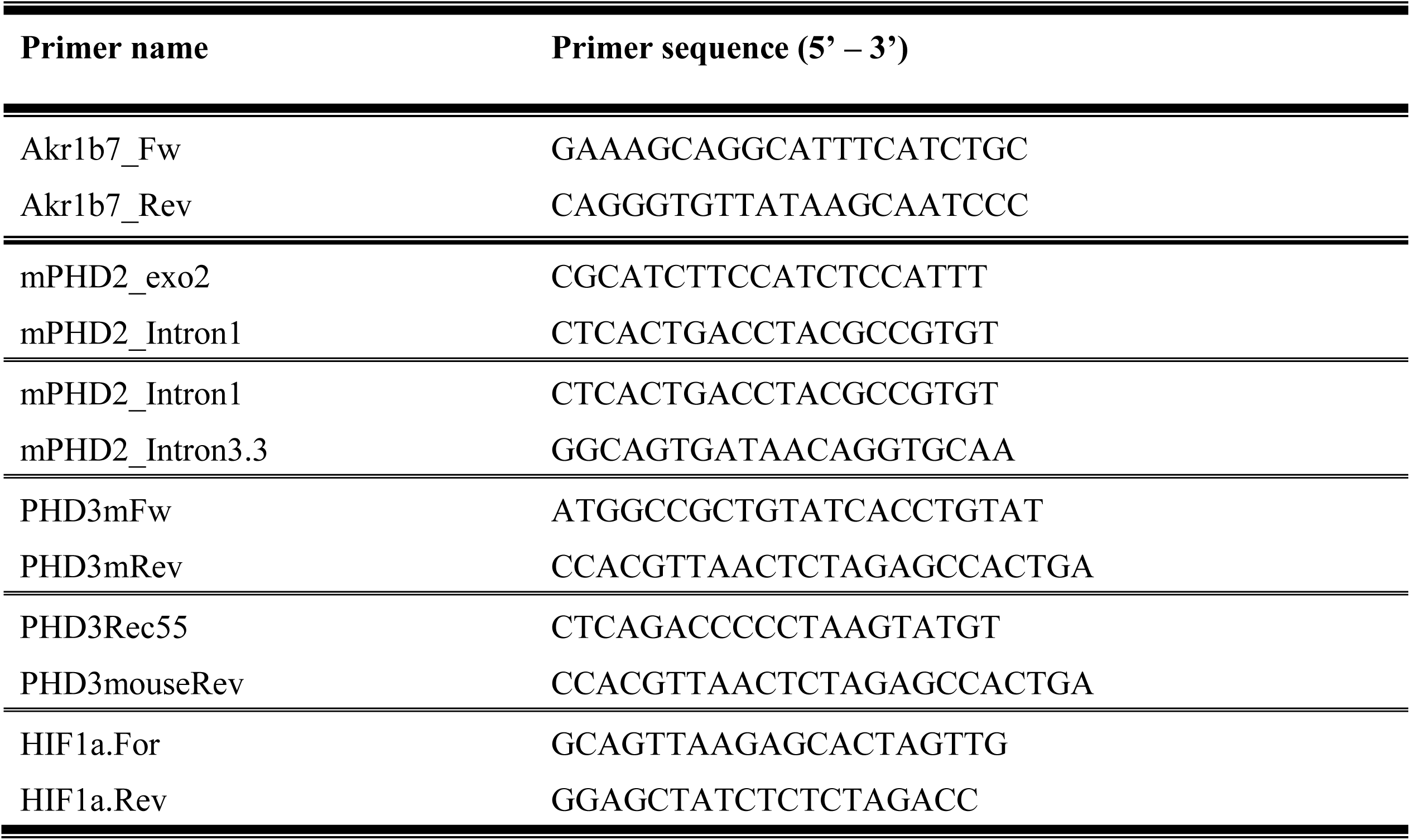
Primers for genotyping of mouse strains

**Table II:**
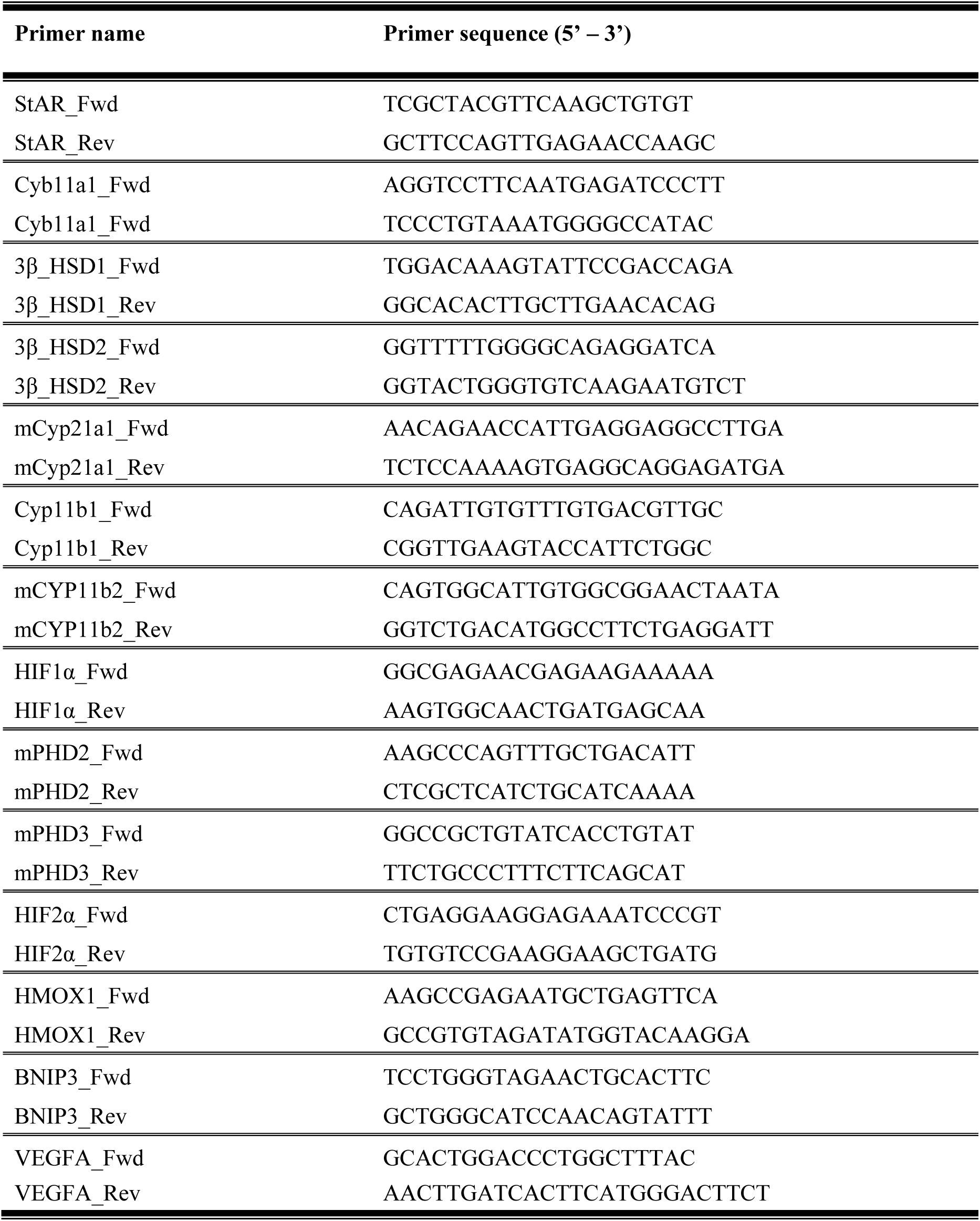
Primers for qPCR analysis

## Notes

### Competing Interest Statement

The authors have declared no competing interest.

